# Stoichiometric balance of protein copy numbers is measurable and functionally significant in a protein-protein interaction network for yeast endocytosis

**DOI:** 10.1101/205674

**Authors:** David O Holland, Margaret E Johnson

## Abstract

Stoichiometric balance, or dosage balance, implies that proteins that are subunits of obligate complexes (e.g. the ribosome) should have copy numbers expressed to match their stoichiometry in that complex. Establishing balance (or imbalance) is an important tool for inferring subunit function and assembly bottlenecks. We show here that these correlations in protein copy numbers can extend beyond complex subunits to larger protein-protein interactions networks (PPIN) involving a range of reversible binding interactions. We develop a simple method for quantifying balance in any interface-resolved PPINs based on network structure and experimentally observed protein copy numbers. By analyzing such a network for the clathrin-mediated endocytosis (CME) system in yeast, we found that the real protein copy numbers were significantly more balanced in relation to their binding partners compared to randomly sampled sets of yeast copy numbers. The observed balance is not perfect, highlighting both under and overexpressed proteins. We evaluate the potential cost and benefits of imbalance using two criteria. First, a potential cost to imbalance is that leftover proteins without remaining functional partners are free to misinteract. We systematically quantify how this misinteraction cost is most dangerous for strong-binding protein interactions and for network topologies observed in biological PPINs. Second, a more direct consequence of imbalance is that the formation of specific functional complexes depends on relative copy numbers. We therefore construct simple kinetic models of two sub-networks in the CME network to assess multi-protein assembly of the ARP2/3 complex and a minimal, nine-protein clathrin-coated vesicle forming module. We find that the observed, imperfectly balanced copy numbers are less effective than balanced copy numbers in producing fast and complete multi-protein assemblies. However, we speculate that strategic imbalance in the vesicle forming module allows cells to tune where endocytosis occurs, providing sensitive control over cargo uptake via clathrin-coated vesicles.

## Introduction

Protein copy numbers in yeast vary from a few to well over a million(1, 2). Expression levels, along with a protein’s binding partners and corresponding affinities, are critical determinants of a protein’s function within the cell. In the context of multiprotein complexes - especially obligate complexes such as the ribosome - it is thought that protein concentrations are balanced according to the stoichiometry of the complex. This is referred to as the dosage balance hypothesis (DBH)(3–5).

For obligate complexes, dosage balance means that there are no leftover subunits, as these would be a waste of cell resources. However, even for proteins in non-obligate complexes a number of deleterious effects could be caused by imbalance. An overexpressed core or “bridge” subunit may sequester periphery subunits, paradoxically lowering the final number of complete complexes(5, 6). Excess proteins may be prone to misinteractions, also called interaction promiscuity, with nonfunctional partners. Numerous studies have identified proteins with high intrinsic disorder as sensitive to overexpression(7–9), and these proteins have low, tightly regulated native expression levels(10, 11) indicating that misinteraction propensity and abundance are related. Underexpression carries its own dangers: a single underexpressed subunit will become a bottleneck for the whole complex. In addition, weakly expressed proteins are noisier(12) and thus less reliable for the cell. Male (XY) animal cells are known to employ “dosage compensation” mechanisms to increase the expression of X-chromosomal genes to be on par with female cells(13, 14), though for other genes it is the female cell that cuts expression levels in half(15), indicating that the cell preserves an optimized set of expression levels.

But optimized does not necessarily mean balanced. Imbalance may be necessary for functional reasons: signaling networks utilize underexpressed hubs to regulate which pathways are active as a given time(16). Recent models show imbalance can be beneficial to complex assembly when affinity and kinetics are taken into account(17, 18). A study of over 5,400 human proteins by Hein et al. found that strong interactions forming stable complexes are correlated with balance, but weak interactions are not, which may mean that the network as a whole is not balanced (19).

Here, we test the hypothesis that protein expression levels are significantly biased towards balance, even for complex PPINs that include weak and transient interactions. This first required us to we develop a method to quantify stoichiometric balance in any arbitrary PPIN, given known binding interfaces and some observed copy numbers. Copy number correlations thus are evaluated beyond direct binding partners to the more global network of interactors. We then can quantify the consequences of imbalance relative to perfect balance according to two criteria: 1) the deleterious consequences and cost of forming misinteractions, and 2) the potentially beneficial control of specific functional outcomes by modulating which complexes, given known binding affinities, actually assemble. Applied to the 56-protein, manually curated, interface-resolved CME PPIN (20), two of its subnetworks, as well as the ErbB PPIN(16), we find that stoichiometric balance in observed copy numbers is often significant, and observed imbalances, particularly of underexpressed proteins, could provide tuning knobs for functional outcomes.

The first consequence of imbalance we evaluate, misinteractions cost, has an indirect effect on function by allowing unbound proteins to bind to non-functional partners, sequestering components and thus affecting formation of specific complexes. They are believed to play a role in dosage sensitivity(7, 8, 21), and avoiding them has been shown to be an evolutionary force limiting protein diversity(22, 23), expression levels(24, 25), binding strengths(26), and protein network structure(23, 27). Misinteractions, not being selected for by evolution, are weak and generally unstable, but there are far more ways for proteins to misinteract than bind to their few functional partners (22, 23). Cells have evolved a variety of mechanisms to increase specificity, such as allostery(28, 29), negative design(30, 31), compartmentalization(22), and temporal regulation of expression(32). Copy number balance would be another such mechanism, as protein binding sites would saturate their stronger-binding functional partners

The second and ultimately more direct consequence of imbalance we evaluate is that changes to copy numbers control which specific and functionally necessary complexes can form. When the central clathrin protein is knocked out in cells, for example, clathrin-mediated endocytosis (CME) is terminated, as clathrin is functionally irreplaceable(33). The plasma membrane lipid PI(4,5)P_2_ is also essential for CME, as it is required for recruiting the diverse cytosolic clathrin-coat proteins to the membrane to assemble vesicles(34). Many clathrin-coat proteins, however, can be knocked out without fully terminating CME(35). As the CME network illustrates (Fig 1), most of these proteins have multiple domains mediating interactions involving both competitive and non-competitive interactions. Adaptor proteins (proteins that bind to the membrane, to transmembrane cargo, and often to clathrin as well) exhibit redundancy in their binding partners that can partially explain how knock-outs to one protein can be rescued by the activity of related proteins. With simulation of simple kinetic models, we can then test these hypotheses. Although these models are far too simple to recapitulate the complexities of CME *in vivo*, they are nonetheless useful in highlighting potential bottlenecks in assembly due to copy numbers or binding affinities.

**Fig 1.**
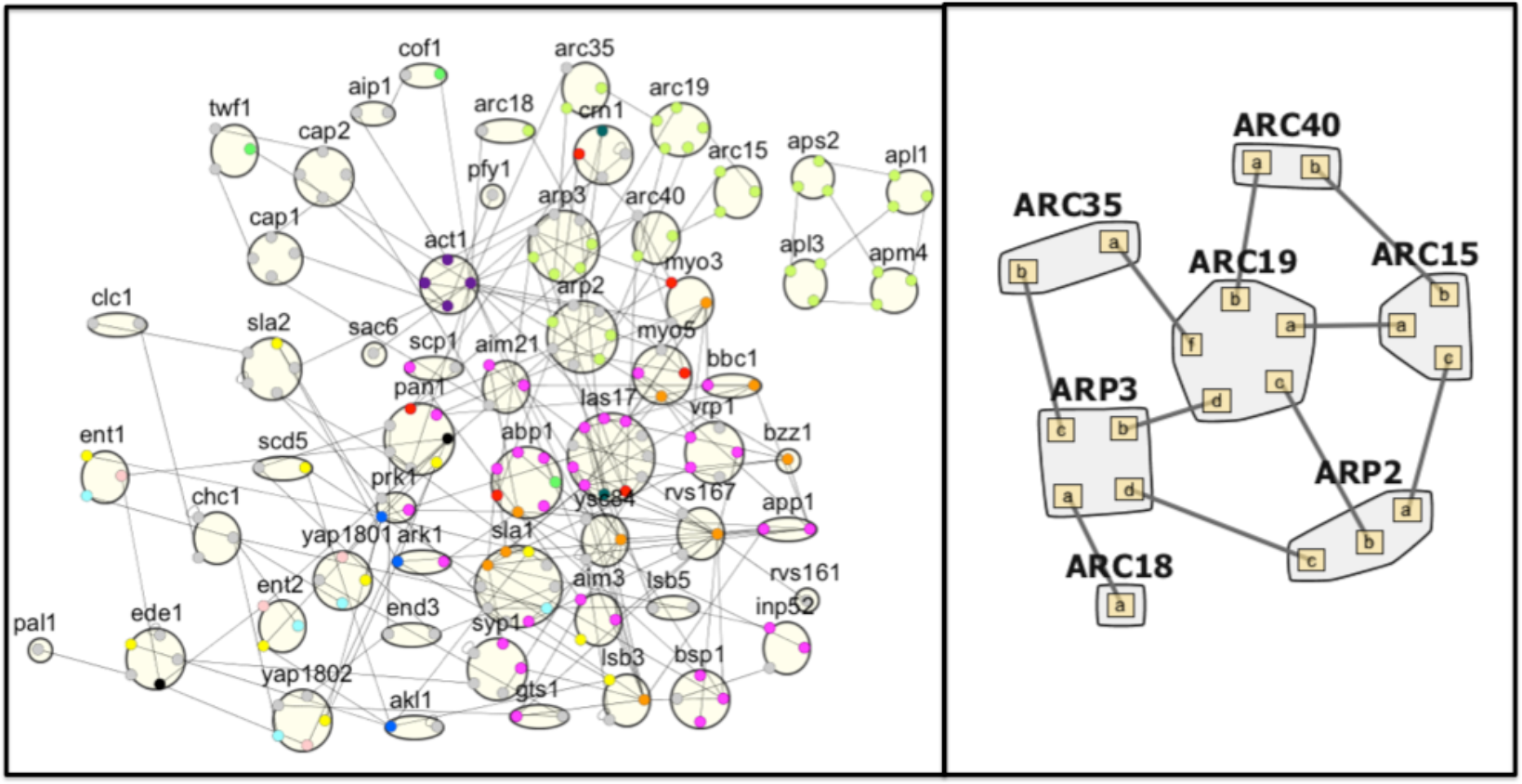
Clathrin-mediated endocytosis network in yeast. (Left) Site graph for the protein-protein interaction network (N=56, E=186), displaying interfaces used for binding interactions. Interfaces are color-coded according to domain type, the most common being SH3 domains (orange), Proline-rich regions (pink), phosphosites (yellow), acidic domains (red), and multi-protein complex subunit interfaces (light green). (Right) The ARP2/3 complex, a subset of the larger CME network.

Quantifying balance in protein networks can thus lead to new insights, as unbalanced proteins may serve as assembly bottlenecks, or maintain alternate cellular functions outside of the network module being analyzed(18). Dosage balance is also important for understanding dosage sensitivity(4, 21), a phenomenon where overexpression of a gene is detrimental or even lethal to cell growth. Studies estimate ~15% of genes in *S. cerevisiae* to be dosage sensitive(9, 36), but the negative effects of gene overexpression have been observed in several eukaryotic species including maize(4), flies(37), and humans(38–40). Studying balance at a network-wide level is challenging because it requires resolved information about the interfaces proteins use to bind. A protein that binds noncompetitively with two partners requires equal abundance to its partners. But if the binding is competitive - i.e. the same interface is used to bind two different partners - the protein’s abundance must equal the sum of that of its partners to have no leftovers. Classic protein-protein interactions networks (PPINs) lack this resolution, but recent studies have begun to add this information, creating what we refer to as interface-interaction networks (IINs)(16, 20, 41). An IIN tracks not just protein partners but also the binding sites that proteins use to bind.

Our study of stoichiometric balance in larger, interface resolved PPINs is organized in three parts. In the first part we define a metric for quantifying stoichiometric balance and how noise in protein expression levels can be approximately accounted for. We apply this to the CME PPIN (20, 41) and the ErbB PPIN (16), highlighting which proteins are over- and underexpressed relative to perfect balance. Although this analysis excludes temporal expression and binding affinity, it provides a starting point for the analysis of these features in the subsequent sections. In part two, we switch to generalized interface-interaction network (IIN) topologies and network motifs to focus exclusively on how our first evaluation criteria, the cost of misinteractions under imbalance, is worse for strong binding proteins and for network topologies that resemble biological networks. In the third part, we return to the interface-resolved CME PPIN to evaluate the observed degree of stoichiometric balance in two smaller sub-networks of the CME network: the 7-subunit ARP2/3 complex and a simplified, nine protein, clathrin-coat forming module. In these sub-modules, we now can also evaluate our second criteria and assess how observed copy numbers influence proper multi-protein assembly given known binding affinities of interactions. Our simulations of (non-spatial) kinetic models demonstrate that stoichiometric balance does, in fact, improve multiprotein assembly relative to observed copy numbers. We speculate that the observed imbalances in clathrin adaptor proteins could offer a mechanism for making the vesicle formation process more tunable, since adaptor proteins are responsible for selecting cargo for endocytic uptake, which is the ultimate purpose of CME.

## Results

### 1. Stoichiometric balance is measureable in large PPINs when interfaces are resolved

For a multi-subunit complex such as the ribosome or ARP2/3 complex (Fig 1b), all subunits bind together non-competitively to assemble a functional complex. Stoichiometric balance is simply having enough of each subunit to form complete complexes, with no subunit in excess. But quantifying balance in a general protein-protein interaction network is more challenging because some proteins will bind competitively, using the same interface for multiple interactions. Such proteins will need a higher concentration in order to saturate their functional partners. Thus, to establish stoichiometric balance in a PPIN the binding interfaces must be known. In previous work we analyzed several interface-resolved PPINs, including the 56-protein clathrin-mediated endocytosis (CME) network in yeast (20, 41) (Fig 1a), and the 127-protein ErbB signaling network in human cells(16).

To balance a network, a number of desired complexes may be assigned to each edge and then the number of required interface copies directly solved for. This is constrained with a starting set of copy numbers, C_0_, otherwise the solution would be arbitrary. However, the inclusion of multiple interfaces per protein introduces a new constraint: interfaces on the same protein should have the same copy number. This constraint often makes nontrivial solutions (i.e. when none of the proteins are set to zero) impossible (see Methods). Therefore, we treat it as a soft constraint, using a parameter “α” to balance its influence. A high α allows more variation of interface copy numbers on the same protein. We constructed and minimized an objective function using quadratic programming (Methods), which produces a new, optimally balanced set of copy numbers, C_balanced_. For any given interface-resolved PPIN, there can be multiple locally optimized solutions of balanced copy numbers.

The benefit of this method is that the distance between C_0_ and C_balanced_ gives you a relative estimate of how “balanced” C_0_ already is, and thus a metric from which to evaluate the significance of balance in the observed copy numbers. Using real copy numbers taken from Kulak et al.(2) (S2 Table), C_real_, as C_0_, we calculated both chi-square distance (CSD) and Jensen-Shannon distance (JSD) between C_real_ and C_balanced_ (Methods). The former metric looks at differences between absolute values and penalizes high deviations more strongly than low deviations, whereas the latter converts both vectors to distributions and measures the similarity between them. We do not expect any networks to have C_real_ that is already perfectly optimized, such that C_real_=C_balanced_. To establish the significance of both distance metrics, we generated 5,000 sets of random C_0_ vectors, sampled from a yeast concentration distribution. We then measured the CSD and JSD from C_0_ to C_balanced_ for each of these random copy number vectors. If C_real_ is balanced, its distance metrics should have a significant p-value relative to yeast copy numbers selected randomly from the yeast distribution.

#### 1.1 Accounting for noise in observed copy number measurements

Even constitutively expressed genes do not have a constant abundance; they vary due to both extrinsic and intrinsic noise (42). Taniguchi et al. found that the abundance of a single protein in *E. coli* follows a gamma distribution (12). Therefore, one reason copy number balance should not be expected to be perfectly matched is due to inherent fluctuations in protein copy numbers. Our algorithm, however, ultimately assigns a single copy number to each interface in the network to optimize perfect balance, when realistically a range of values would be more appropriate.

Our method does provide one mechanism to allow a range of copy number values for a single protein, and that is through allowing each interface on a single protein to have distinct values. This range can be tuned through our parameter α, which biases solutions towards equivalent interface copies per protein when set to zero. As the α parameter increases, more variation is observed. For example, one interface may be assigned 200 copies and another on the same protein 300 copies. If the protein is usually expressed within the 150-350 copy range, this solution is more realistic than enforcing both copy numbers to be exactly 250.

We therefore systematically characterized how variations in α changed the “noise”, or variability in interface copy numbers on each protein. Taniguchi et al. found that yeast proteins with high abundance (~1,000 or more copies) had a noise (σ^2^/μ^2^) upper limit of about 0.5 with ungated data and 0.1 with gated data(12). For α≤0.03, we found that proteins with mean interface copy numbers above 1,000 had less than 0.1 noise, indicating that such a solution is possible. (S1 Fig). Low abundance proteins exhibit higher noise in terms of expression level(12, 43), and this feature is also observed in our model. We therefore used values of α in the 0.01 to 2 range based on this analysis (S1 Fig).

#### 1.2 Protein copy numbers in yeast clathrin-mediated endocytosis are balanced

As Fig 2a,b shows, at α=1 the p-value for JSD was found to be statistically significant (p=0.0054) but the p-value for chi-square distance was not (p=0.157). We analyzed the real copy numbers before and after balancing and found that the protein cofilin was highly overexpressed (Fig 2c) meaning that it had to be greatly lowered to achieve balance. This resulted in a skewed CSD for C_real_, which the change in cofilin dominated. We therefore re-tested the degree of balance when cofilin was removed from the network. At α=1, both JSD (p=0.0012) and CSD (p=0.022) were statistically significant (Fig 2d), indicating that these 55 proteins are balanced compared to random copy numbers. These results were robust to changes in α, but the p-values tended to be lowest when α was in the 0.01 to 2 range. The absolute distance from C_real_ to C_new_ lowered as α was raised, plateauing when α≥10.

**Fig 2.**
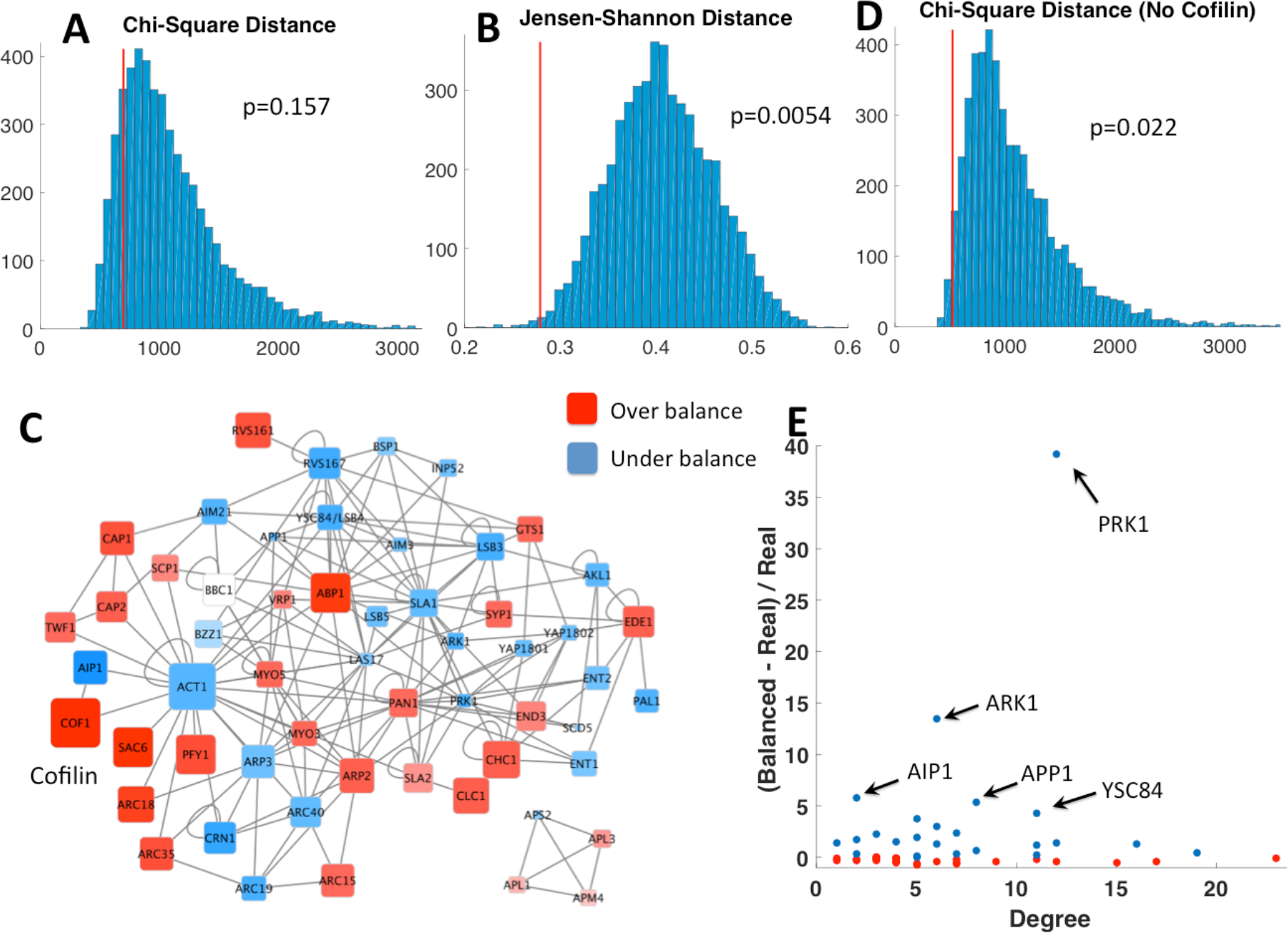
Clathrin-mediated endocytosis proteins are balanced. **(A,B)** Histograms for chi-square distance and Jensen-Shannon distance between the real protein copy numbers and their copy numbers after balancing. Compared to 5,ooo sets of random sampled copy numbers, the real copy numbers had a statistically significant Jensen-Shannon distance, but not chi-square distance. **(C)** Graph of CME network, showing which proteins were overexpressed (red) or underexpressed (blue) compared to the balanced copy numbers. Cofilin was highly overexpressed, which led to a high chi-square distance. **(D)** Histogram for chi-square distance when cofilin was removed from the network. It is now statistically significant, indicating that theother 55 proteins are balanced compared to random copy numbers. **(E)** The five most underexpressed proteins were two kinases (PRK1 and ARK1), one phosphatase (APP1), and two partners of Actin (AIP1 and YSC84). The former three bind transiently to their partners, so there is no functional need for them to be balanced. The latter two are discussed in the text.

Because protein complexes that strongly bind are thought to be more balanced than weak interactions, we repeated the analysis on the full 56-protein network after removing one of two modules from the network: the four protein subunits of the AP complex, and the seven proteins in the ARP2/3 complex. Without the former, the p-value increased to 0.0088 for JSD and 0.197 for CSD, indicating less overall balance. Removing only the ARP2/3 complex similarly raised the p-values to 0. 023 and 0.24. This trend held when cofilin was also removed.

The four AP subunits that form the obligate AP-2 complex are fairly close in abundance, as are the clathrin heavy chain and clathrin light chain proteins, which is consistent with the pressure for strong binding proteins to be more tightly balanced.

#### 1.3 Stoichiometric balance is not measured without proper interface binding interactions

To test whether balance depended mostly on protein network structure rather than the child interface interaction network (IIN) structure, we ran this analysis again using random IINs for the same parent protein network, again excluding cofilin. In other words, we randomized whether proteins bind competitively or noncompetitively, using a rewiring method from Holland et al.(41). For 20 random IINs, we found that the real copy numbers were significantly less balanced. For α=1, the same analysis obtained p-values of 0.44 ± 0.12 for CSD and 0.24 ± 0.13 for JSD. Thus the protein copy numbers are balanced according to the underlying interface network.

#### 1.4 Observed protein imbalances can highlight functional relationships

Finally, by looking at the relative change between C_real_ and C_balanced_, we could examine which proteins are underexpressed in the network. As Fig 2e shows, the five most underexpressed proteins are PRK1 (by a factor of nearly 40), ARK1, AIP1, APP1, and YSC84. PRK1 and ARK1 are both kinases; they form transient interactions which their partners for the purpose of phosphorylation. Since a single kinase can phosphorylate many proteins relatively quickly, rather than form stable complexes with each target, there is a sensible functional explanation for why these proteins can be underexpressed relative to their partners by such a large margin. Similarly, APP1 is a phosphatase. The protein AIP1 is an actin binding protein that targets a binding surface of actin without any competition from other actin binders, and also binds the highly expressed cofilin. Its low abundance relative to actin and cofilin could indicate it acts as a bottleneck in regulating cofilin-actin interactions, or perhaps more simply, that functionally it is not needed at a 1:1 stoichiometry with the ubiquitous actin protein. YSC84 has 13 binding partners, and 10 of these partners all bind the YSC84 SH3 domain, including the relatively highly expressed ABP1. Although many of these binding partners (all proline rich domains-PRDs) also have additional partners of their own, ABP1’s PRD is specific to YSC84’s SH3 domain(41). As we return to in the discussion, underexpression could indicate a functional regulatory role for this protein, or indicate transient interactions with partners. Identifying underexpressed proteins and which of their interface binding partners apply pressure to increase copy numbers is a useful first step in hypothesizing about the temporal dynamics of such proteins within the cell.

#### 1.5 Upstream proteins in the ErbB signaling network are underexpressed

We applied our algorithm to another IIN from the literature: that of the 127 protein human ErbB signaling network, characterized by Kiel et al.(16). Our algorithm optimizes copy numbers to the full network structure even if not all individual target copy numbers are available. Thus we measured the distance between the real (C_real_) and optimized (C_balanced_) copy numbers for the 115 of the 127 proteins for which we could assign expression levels from HeLa cells (Methods). We compared results to copy numbers randomly sampled from a HeLa protein concentration distribution.

Because this is a signaling network where the majority of interactions are phosphorylation, we expected these transient interactions to bias the copy numbers against significant balance. However, while JSD was not found to be significant (p=0.274), CSD was (p=0.022). This result held when copy numbers were shuffled rather than randomly sampled (JSD: p=0.120, CSD: p=0.019). As stated above, CSD is dominated by large deviations. Thus, while the network as a whole is not balanced, there appears to be no dramatic overexpression.

The three Ras proteins (HRAS, NRAS, and KRAS) were found to be underexpressed (S2 Fig), confirming the findings of Kiel et al. using simpler comparisons of Ras copy numbers to all binding partners (16). Also found to be underexpressed were all five MAP3K proteins (RAF1, MAP3K1, MAP3K11, MAP3K2, and MAP3K4) in the network. MAP3K proteins are the top layer in MAPK cascades, a signaling motif consisting of three proteins (a MAP3K, MAP2K, and MAPK) occasionally bound together via a scaffold protein(44). The membrane bound receptors ErbB2 and ErbB3 were similarly underexpressed. These proteins are all “upstream” in the ErbB signaling network. These conserved correlations in copy numbers suggest an interesting trend; protein abundance is limiting at the start of a signal cascade, potentially to control specific outputs from diverse inputs, and increases as a signal travels “downstream” in a signaling network.

### 2. Imbalance increases misinteractions dependent on the network topology and binding affinities of proteins

In this second part, we investigate how the cost of imbalance, measured solely in terms of misinteractions, depends on general properties of proteins, including binding affinity and number of binary partners. In a stoichiometrically balanced network, proteins will be driven to saturate their stronger-binding functional partners. Any “leftover” proteins, however, may misinteract, or form non-functional complexes that, while weak, are combinatorially numerous.

#### 2.1 Misinteractions are minimized under balanced copy numbers and are largely independent of network motif structure

Complex formation and misinteractions must be evaluated at the level of individual protein binding interfaces, and we thus study small network motifs that have been previously characterized in real biological interface interaction networks (IINs) to control binding specificity (41). Of these five motifs (Fig 3a), the hub and square motif are the most common in biological IINs relative to random networks (41). The chain, triangle, and flag motif are selected against due to the challenges in optimizing such binding interfaces for strong selective binding and against misinteractions.(23, 27, 41) The motif defines the functional or “specific” interactions, which we allow at equal binding strengths. However, all other possible protein-protein interactions were allowed as misinteractions, which occur at weaker strength than the specific interactions. Because each node represents an interface (each on its own protein in this case), all binding was competitive.

**Fig 3.**
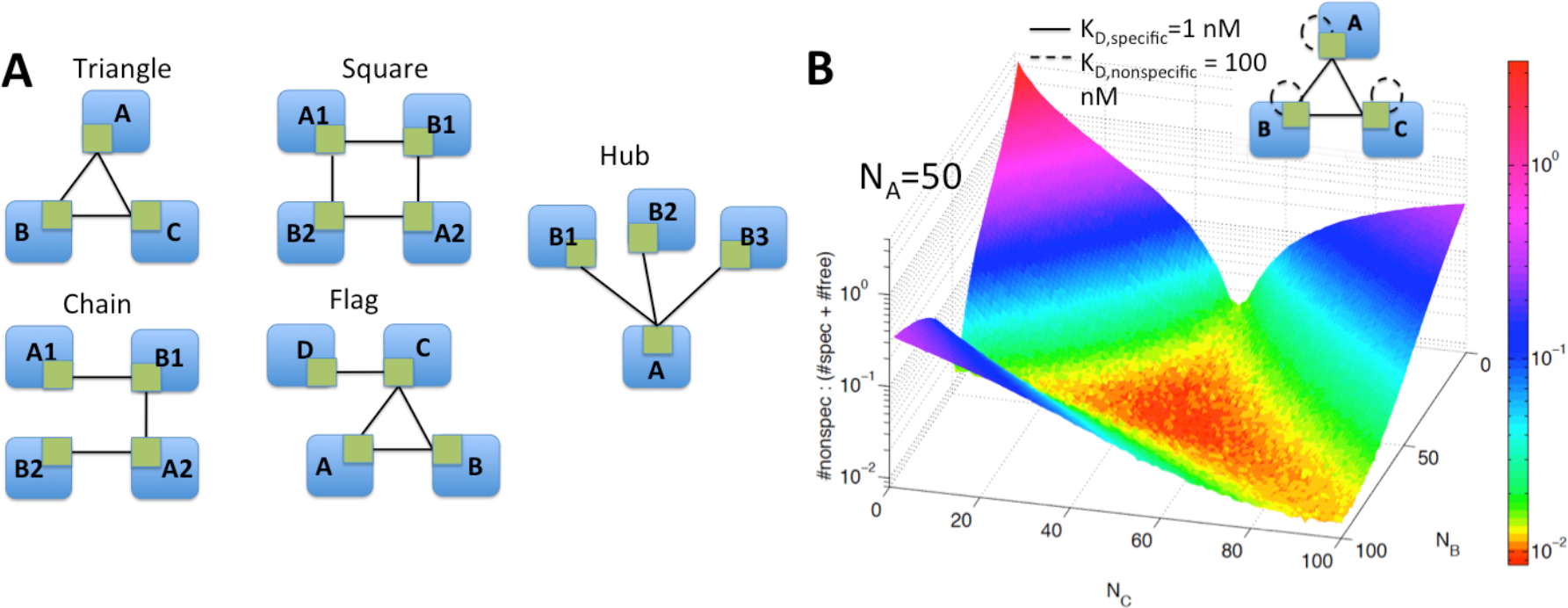
Misinteractions in network motifs from biological IINs. **(A)** Five network motifs that have been shown to impact specificity of binding in biological IINs were tested for the effects of imbalance on misinteractions. **(B)** Surface plot obtained for the triangle network. The z-axis is the frequency of misinteractions at steady-state (Cost: Eq. 1) averaged across 1000 runs. The x and y axes are the number of B and C proteins; the number of A proteins is fixed at 50. As one protein becomes overexpressed, misinteractions increase exponentially.

Balanced copy numbers are relatively easy to design for these simple network motifs, and the optimization of the previous section is not necessary. We study imbalanced copy numbers by simply varying the copy numbers of two proteins in each network over a wide range while keeping the remaining proteins constant. For each set of copy numbers, we ran the system to equilibrium using the Gillespie algorithm(45). We could then measure the total number of specific and non-specific complexes formed (N_specific_, N_nonspecific_), as well as unbound proteins (N_free_), and use this to evaluate the cost of being out-of-balance in terms of misinteraction frequency:

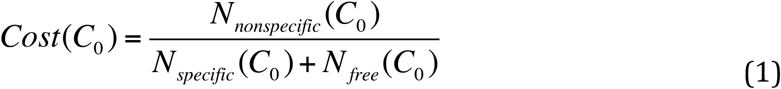

averaged across 1,000 runs, where C_0_ is the vector of initial copy numbers.

The frequency of misinteractions is lowest when the protein copy numbers are balanced. Fig 3b shows the results for the triangle network. For example, when all three proteins have equal abundance of 50 copies, about 25 of each specific complex are formed, and minimal proteins are leftover. Cost also remains low when two proteins are equally overexpressed, as these excess proteins can bind to each other. The instances where misinteractions are the most frequent are when one protein is overexpressed, as this protein has no specific partners left and thus will self-bind: a misinteraction for this motif. Similar surface plots were obtained for all five network motifs (S3 Fig).

Notably, with balanced copy numbers, the frequency of misinteractions is almost entirely dependent on the relative strength, or energy gap, between specific and nonspecific binding (Fig 4a) and there was little difference among the five networks. The slope varies slightly from one motif to another, and we confirmed that this can be calculated relatively accurately based on the ratio of specific versus non-specific interactions possible for that motif. Furthermore, the results were similar when we varied the absolute strength of specific binding from 1nM; under balanced conditions it affects the number of free proteins (N_free_) relative to total complexes formed. Thus, under balanced copy numbers, the cost of misinteractions is not strongly dependent on specific binding affinities.

**Fig 4.**
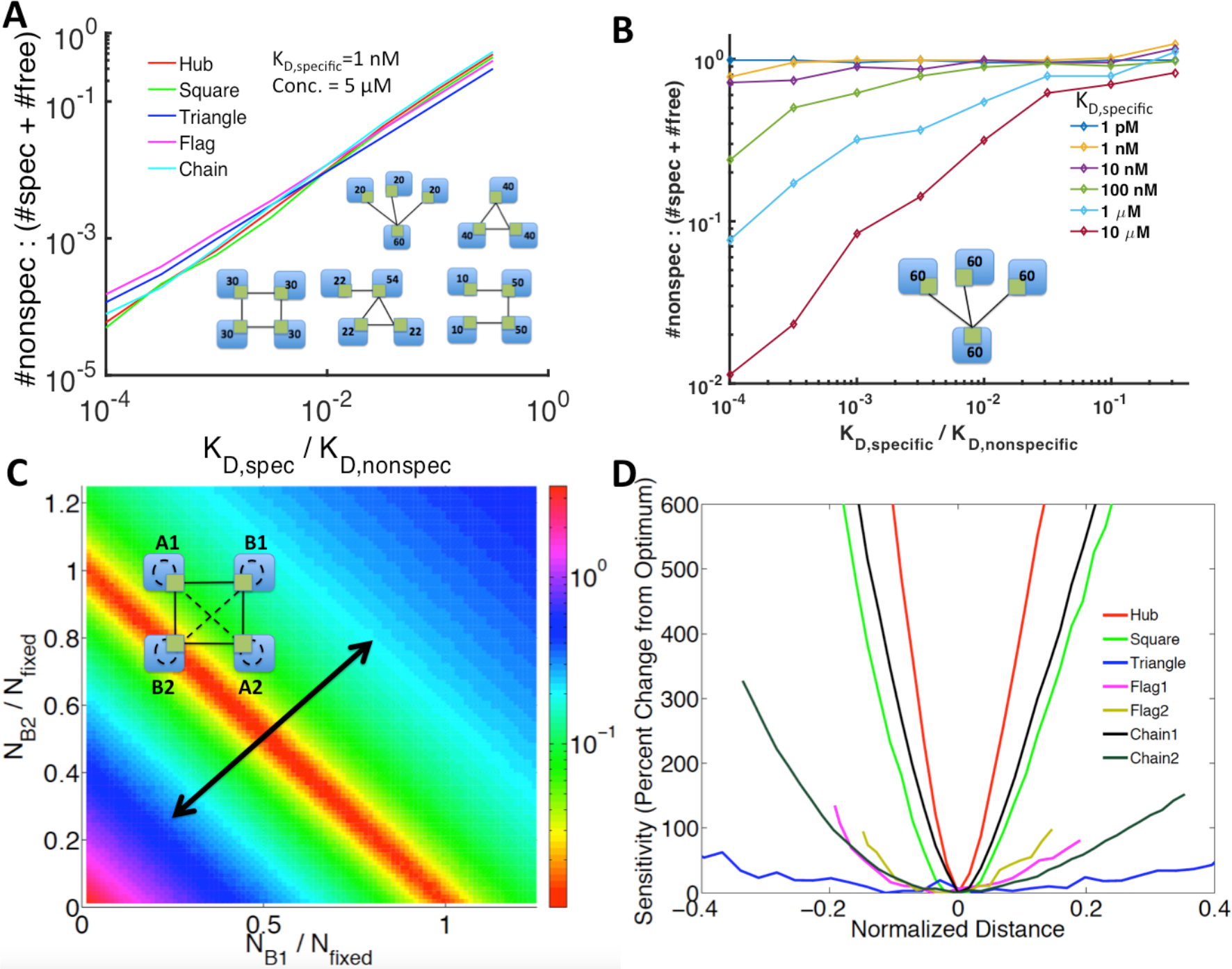
Misinteractions are motif dependent only when concentrations are imbalanced. **(A)** At balanced concentrations, misinteraction frequency increased linearly with the ratio of K_D,specific_ to K_D, nonspecific_. It was also roughly equal for all five network motifs. **(B)** At unbalanced concentrations, misinteractions can occur even at a low energy gap, unless the overall binding is weak. **(C)** Surface plot for the square network, measuring the ratio of (#nonspecific complexes: #specific complexes + free proteins) when A1 and A2 are fixed while B1 and B2 are varied. The principal component (black line) is shown across the region of lowest misinteraction frequency. **(D)** Cost sensitivity to concentration imbalance varies significantly between motifs. The “distance” is measured along the principal component of the surface plots as you move away from the optimal region. Two different pairs of fixed proteins were analyzed for the chain and flag networks. The hub and square networks were the most sensitive to imbalance, while the flag and triangle were the least.

#### 2.2 Misinteractions for imbalanced copy-numbers are worse for biologically common motifs and strong binding proteins

Unlike the similar cost of misinteractions under balanced copy numbers, the five networks noticeably differ in sensitivity to imbalanced copy numbers. In general, as copy numbers become more imbalanced, the misinteraction cost grows. To quantify this rate for each network motif, we measured the percent change in cost as one travels along the principal components away from the balanced copy numbers (Fig 4c; S3 Fig). The hub and square motifs were found to be the most sensitive, showing a rapid increase in cost of misinteractions as imbalance grows, whereas the flag and triangle motifs were found to be the least. (Fig 4d). The triangle motif has the least sensitivity and it also has the fewest misinteractions possible; it can form 3 specific complexes and only 3 misinteracting complexes. The robustness of this module also then extends to the flag motif, which contains a triangle.

The motifs most sensitive to imbalance, the hub and square motif, are also the motifs most common in biological networks (41). In previous work, we demonstrated that these motifs are evolutionarily selected for in biological networks because binding interfaces that interact through these specific motifs are much easier to simultaneously design for high specificity (strong K_D,specific_) and for weak nonfunctional interactions (weak K_D,nonspecific_) (41). Although these motifs thus produce more selective binding interfaces, our results show that there is more pressure to maintain copy number balance in these biologically common motifs to prevent misinteractions.

Importantly, unlike the results for balanced copy numbers, strong binding proteins are highly prone to misinteractions under imbalanced conditions (Fig 4b). Weak-binding proteins form minimal complexes overall, and thus imbalances in copy numbers do not strongly influence their binding patterns. Strong binding proteins, on the other hand, are driven to bind to any unbound interface, even when the gap separating specific and non-specific binding is high, and thus leftover copies of these proteins frequently misinteract. This supports the observations that strong binding proteins should be tightly regulated to maintain stoichiometric balance(19), and therefore avoid misinteractions. For weak binding proteins, on the other hand, misinteraction cost is not a significant pressure favoring copy number balance.

#### 2.3 Larger networks with biological topologies produce more misinteractions under copy number imbalance

Our analysis of network motifs above demonstrated that topologies common in biological IINs are actually more prone to misinteractions when copy numbers are imbalanced. We find here that the same trend applies to much larger networks that again exhibit biological topologies (Fig 5). To show this, we analyzed 500 IINs that differed in three properties: motif frequencies; degree distribution; and density, which was determined by the size of the network (90-200 proteins for 150 edges). The biological-like IINs have motif frequencies biased to hub and square motifs, they have a degree distribution that is power-law like or “scale-free”, meaning, broadly speaking, that a few “hub” proteins have many connections while the majority are specialized for a few interactions, and they tend to be sparse; interfaces in the CME IIN have an average degree of only 2.06(41). For simplicity, here we will assume each interface is on its own protein, such that the PPIN is the same as the IIN. Balanced copy numbers are assigned to each network using our optimization method described above based on network structure (also see Methods), and imbalanced copy numbers are defined by randomly sampling copy numbers from the yeast distribution. Specific and non-specific K_d_ values for each possible binding interaction were initially taken from a previous study(27), where the gap between specific and non-specific binding was optimized based on selecting amino-acid sequences for each interface (27).

**Fig 5.**
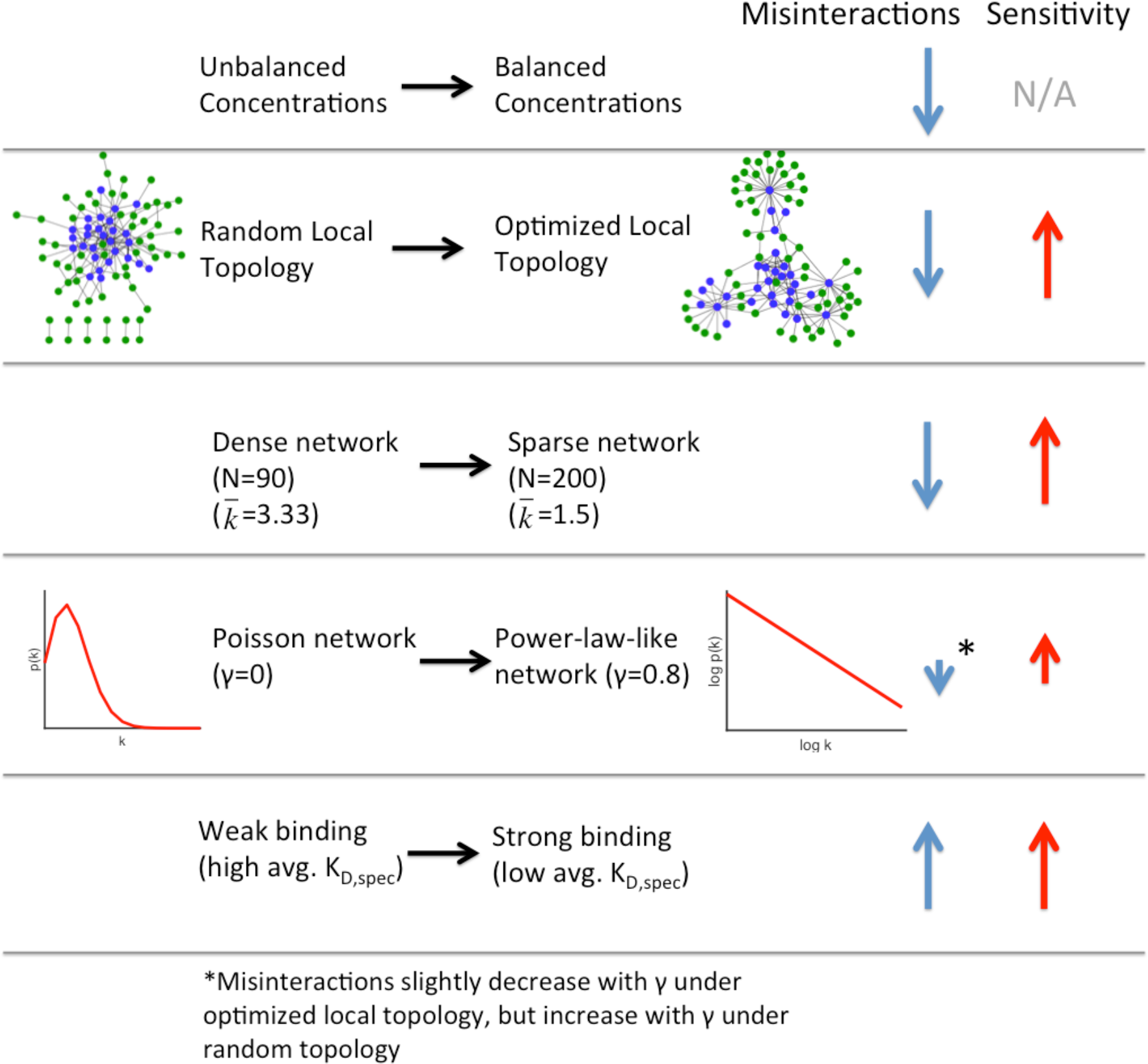
Biological IIN topologies have more misinteractions under imbalance. Shown are trends in misinteraction frequency under balanced concentrations (blue arrows) and sensitivity to imbalance (red arrows). Several features that make networks perform better under balanced concentrations make them perform worse under unbalanced concentrations: sparseness, a topology that matches with real interface networks, and a power-law degree distribution. Strong average binding caused both increased misinteractions and increased sensitivity.

As expected, when copy numbers are balanced rather than imbalanced via random assignments, all networks produced fewer misinteractions. The networks that, under balanced copy numbers, produced the fewest misinteractions were the networks most like biological IINs: they were sparse networks and they had optimized topologies favoring square and hub motifs (Fig 5). Because these IINs also had larger energy gaps separating K_D,Specific_ and K_D,Nonspecific_ (27), we verified that when all networks were assigned the same K_D,Specific_ and K_D,Nonspecific_ (1000-fold different), the biological IINs indeed produced fewer misinteractions under balanced copy numbers (S4 Fig), although the difference was relatively small. Hence, overall, the results are similar to the findings with motifs, that for balanced copy numbers, misinteractions are not strongly influenced by network structure.

Once copy numbers were imbalanced, however, the biological-like IINs produced a sharper increase in misinteractions (higher sensitivity-Fig 5). This is consistent with the trends from the previous section, where the biological motifs of hub and square motifs were also more sensitive to imbalance. Sparse networks are more sensitive to imbalance because they have more interfaces (N) that can possibly misinteract (order N^2^). The only network feature that did not have a significant trend in controlling misinteractions either for balanced or unbalanced copy numbers was the degree-distribution. For power-law network topologies compared to Poisson networks, misinteractions could be higher or lower depending on the local motifs or the network sparseness (Fig 5; S4 Fig). Thus local topology and density was more important than the overall degree distribution.

Finally, because highly abundant proteins are thought to have low average affinity to avoid misinteractions, we increased the absolute strength of K_D,Specific_, while keeping the gap between K_D,Specific_ and K_D,Nonspecific_ the same. Stronger affinity did indeed lead to both more nonspecific complexes and higher sensitivity to copy number imbalance. This result is consistent with the previous section and confirms that strong binding affinities can be paradoxically deleterious to specific complex formation.

### 3. Beyond misinteractions: Multi-protein functional assemblies are sensitive to stoichiometric balance

In the above section we only studied binary, competitive interactions. But proteins often bind noncompetitively into higher complexes, and they may interact weakly and thus form few complexes, in which case imbalance may have functional benefits(17, 18). We created kinetic models of two modules from the CME network with observed imbalances: the ARP2/3 complex and a simplified vesicle forming protein subset. Simulating higher complex formation is challenging because of the exponentially large number of possible species, so we used NFSim(46), a stochastic solver of chemical kinetics that is rule-based, enabling an efficient tracking of higher-order complexes as they appear in time.

#### 3.1 The ARP2/3 Complex has higher yield under stoichiometric balance

One unexpected imbalance we found was that of the isolated, 7-component ARP2/3 complex. The complex has one highly underexpressed subunit, ARC19. ARC19 is a core subunit, binding to five other subunits (Fig 1b). Because of this, it is more likely to form misinteractions (due to its five interfaces) and be a part of incorrect complexes (e.g. complexes of the form ARC19 - ARC40 - ARP2 - ARC19 are incorrect because they contain two ARP19 proteins). Thus we tested whether the observed copy numbers might improve formation of complete ARP2/3 complexes.

Ultimately, we found that balanced copy numbers always improved formation of complete ARP2/3 complexes relative to the observed copy numbers, whether or not misinteractions were modeled (Fig S5). We simulated simplified complex assembly using arbitrary rate constants and two sets of copy numbers: those observed from Kulak et al. and stoichiometrically balanced (in this case equal) copy numbers for each subunit. We measured “yield” as the number of proteins in full complexes divided by the number of proteins in all complexes, including misassembled or incomplete. Some cooperatively was allowed in that if three proteins in a trimer were held together by two binding events, the third binding event could occur at a faster rate (due to all three subunits being localized together). Binding to the core subunit ARC19 was also set to be 10-fold stronger than peripheral bindings, as this increased yield. But no matter what parameter ranges we used, we could not increase the yield of the Kulak copy numbers (max ~13%) versus the balanced copy numbers (max ~50%). Because ARC19 has ~5-fold underexpression compared to the other 6 subunits, incomplete complexes dominate. The results held when we also allowed ARC19 to form misinteractions.

Imbalances in copy numbers have been shown to actually improve the yield for self-assembly, but the optimal copy numbers must take on specific ratios of components to optimize yield (17). Here, we see that the ARP2/3 subunits do not exhibit optimal expression for yield in our model. One possible explanation is that the ARC19 subunit has distinct thermodynamics or kinetics that are critical for controlling assembly. This would suggest that this subunit has conserved expression across all organisms. However, this is not the case. We compared the expression levels of the seven subunits with data from three other studies. Two also found ARC19 to be underexpressed(47, 48), whereas one(1) found it to be overexpressed. However, Chong et al. also found ARP2 to be underexpressed, whereas Kulak et al. found it to be overexpressed. We also compared the abundance of human homologs from five studies(2, 19, 49–51) and found similar issues with noise, though only one found ARC19’s homolog to be underexpressed. (S5 Fig) Thus, no conservation of subunit expression levels is observed. Without a more structurally and biochemically accurate model for the ARP2/3 components, it is difficult to assess whether the low expression of ARC19 does provide some benefit in assembly yield. As we return to in the discussion, several other factors may explain the imbalance, such as noise in expression levels or in measurements of expression levels, or additional roles in the cell for some ARP2/3 subunits.

#### 3.2 A simplified clathrin-coated vesicle forming module enables a kinetic study of imbalance effects on non-equilibrium assembly

For our final analysis we test the effects of copy number balance on a more complex, non-equilibrium model of clathrin-coat assembly for vesicle formation. Our minimal model for vesicle formation includes nine cytoplasmic proteins plus the plasma membrane lipid recruiter PI(4,5)P_2_, with the biochemical parameters taken from the literature for all known binding interface interactions (Fig 6; Table 1). In clathrin-mediated endocytosis, clathrin triskelia consisting of three heavy chains (CHC1) and three lights chains (CLC1) are recruited to the membrane via adaptor proteins that bind lipids (ENT1 & 2, SYP1, SLA2, YAP1801) and in some cases also transmembrane cargo (ENT1 & 2, YAP1801). Clathrin polymerize to form a hexagonal clathrin cage of ~100 triskelia (52) that helps deform the plasma membrane into spherical membrane vesicles of ~100 nm in diameter. Additional non-membrane-binding scaffold proteins help stabilize the assembly (EDE1, YAP1802). Importantly, the assemblies do not have to exhibit a perfect stoichiometry of components, unlike the ARP2/3 complex, in order to function, with variable compositions shown to produce clathrin-coated structures *in vitro*{Dannhauser, 2012 #259;Mishra, 2002 #260;Kelly, 2014 #253}. To measure vesicle formation in our model, we therefore make the assumption that completed vesicles contain 100 triskelia (52) in a complex on the membrane. Once a completed model vesicle is formed, all components that are a part of this complex are recycled, unbound, back to the cytoplasm, keeping total protein concentrations fixed.

**Fig 6.**
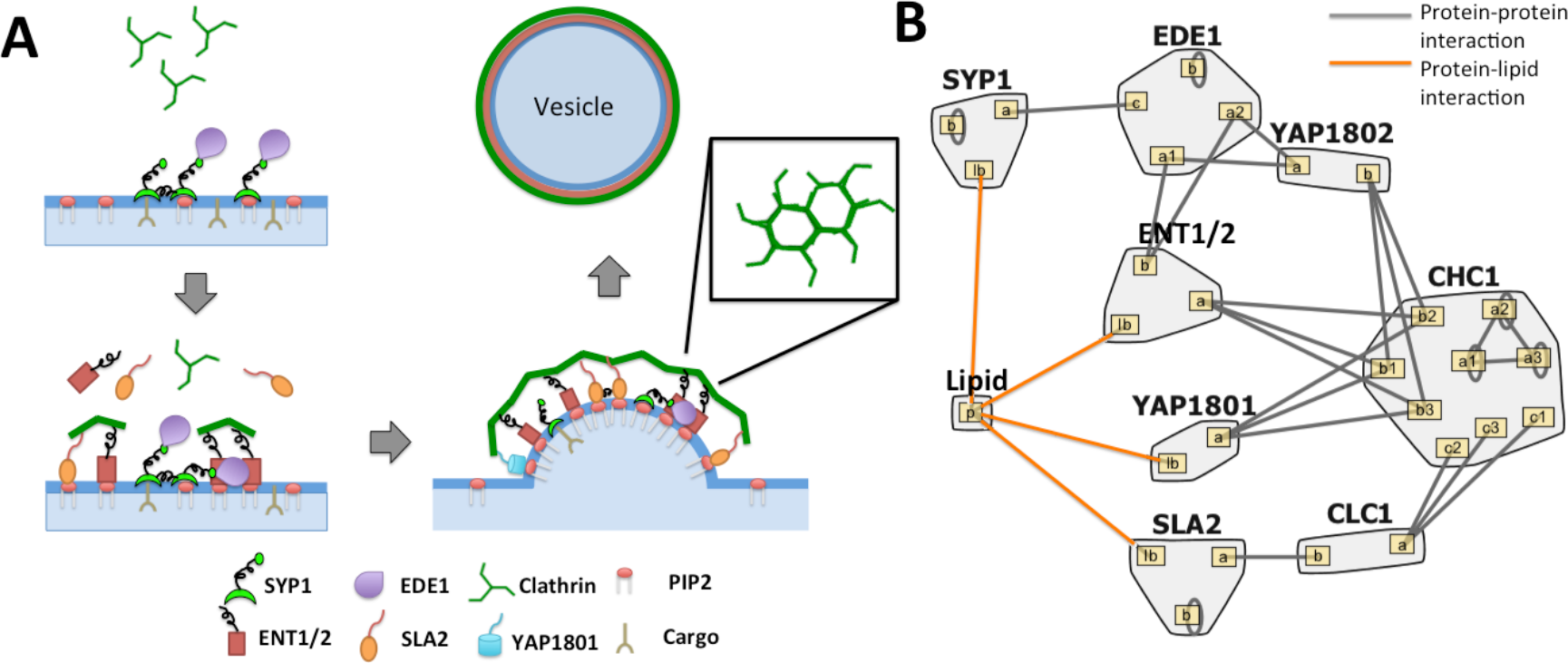
Clathrin membrane recruitment model. **(A)** In clathrin-mediated endocytosis, adaptor proteins bind to the lipid membrane and recruit clathrin triskelia to the surface. These triskelia assemble a hexagonal cage around the plasma membrane vesicle. **(B)** Binding model of the clathrin module. Included are seven adaptor or accessory proteins (SYP1, EDE1, YAP1801/2, ENT1/2, and SLA2), clathrin heavy chains already assumed to be in trimer form, and clathrin light chains. Five of the adaptor/accessory proteins can bind directly to the lipid membrane. Picture generated with Rulebender.

**Table 1:**
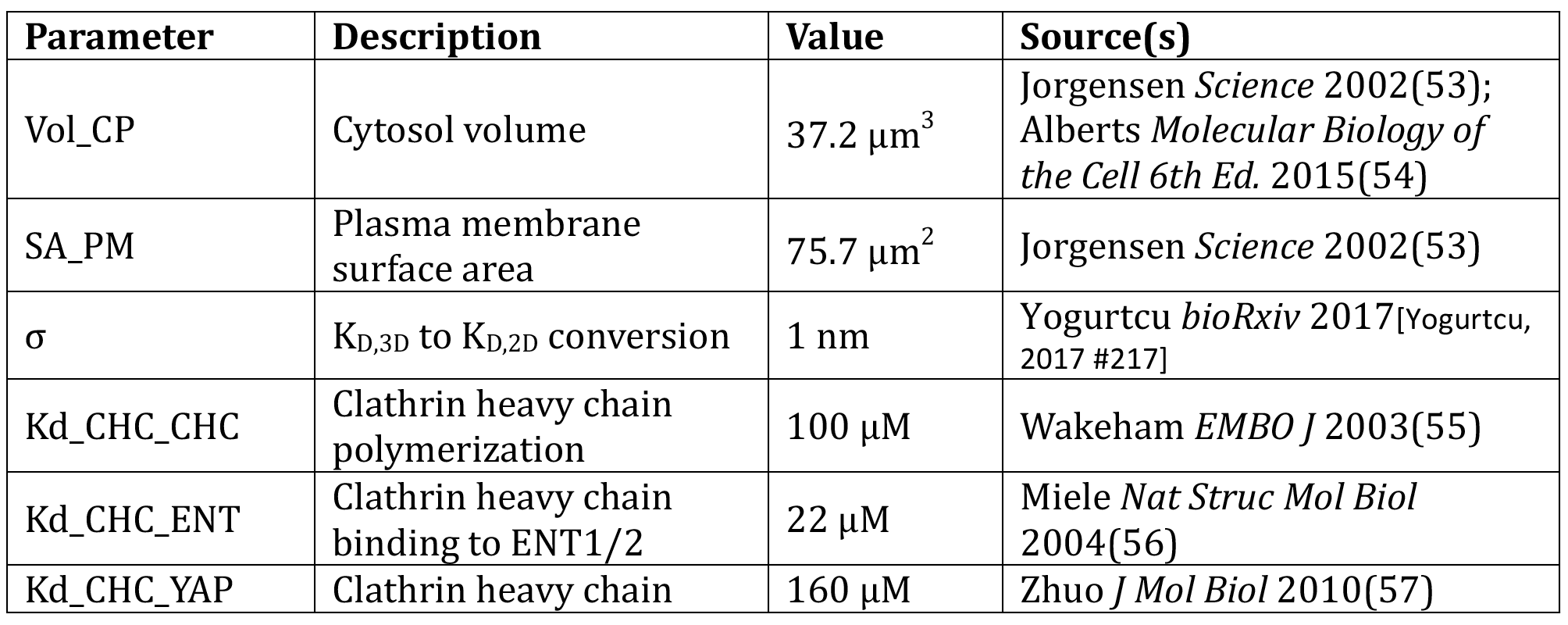
Parameters for clathrin membrane recruitment model. See SI for further notes.

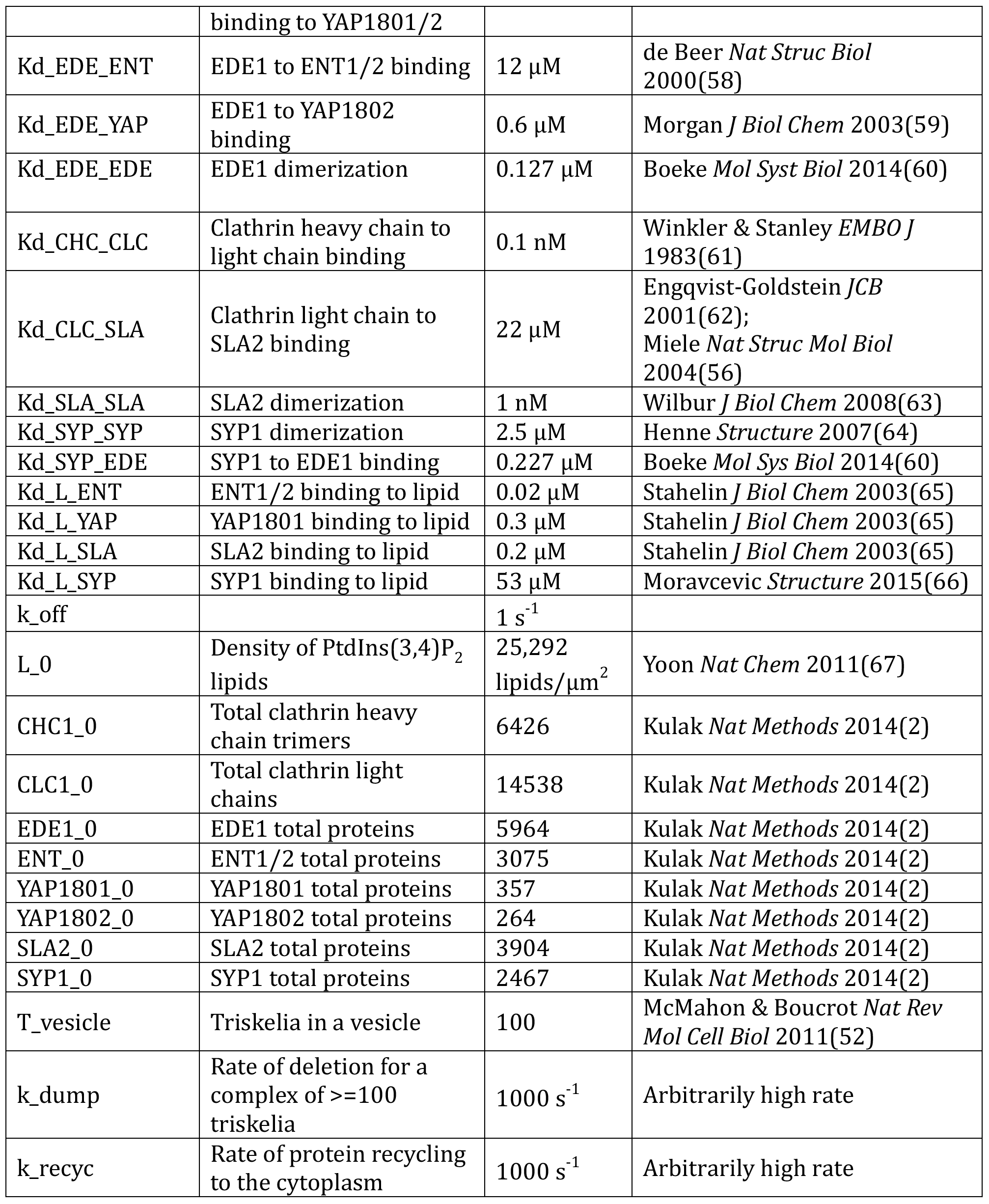

We emphasize that this minimal model is based on the known concentrations and binding properties of the component proteins, and thus we are not attempting to optimize the model to best describe *in vivo* observations. Furthermore, this kinetic model does not account for biomechanics of the membrane budding or coupling to the cytoskeleton, or molecular structure, which are important features of CME. As we see in our simulations, our vesicles form ~10 times faster than vesicle formation *in vivo*. However, clathrin-coated vesicles (pre-scission) are observed to assemble *in vitro* with minimal components, without the cytoskeleton or any energy sources(35, 68). We thus included in our model all proteins from the larger CME network (Fig 1) that directly connect clathrin coat assembly to the membrane surface, linking the assembly process with the ultimate endocytic goal of transmembrane receptor and cargo uptake. Our model thus represents a useful qualitative framework to assess how stoichiometric balance in clathrin-coat components can impact vesicle formation and thus cargo uptake.

An important feature that our model does capture is the reduction in dimensionality (3D to 2D) which accompanies binding to the membrane surface (69). Once localized to the membrane via either lipid binding or recruitment by other proteins, proteins are concentrated in units of Area^−1^, with binding constants of 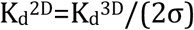, where σ is a lengthscale in the nanometer range{Wu, 2011 #256}, as discussed in Ref. (69). Transitioning to the membrane can drive dramatic increases in complex formation due to higher effective concentrations of components (69). In our simulations here, we find that this is a critical factor controlling vesicle formation. Besides this division between the cytoplasm and the membrane surface, there is no other spatial resolution. A full list of model assumptions can be found in the Supplemental Text.

#### 3.3 Adaptor proteins are underexpressed and can tune vesicle formation

We first evaluated whether this nine-protein module (Fig 6) was significantly balanced. The clathrin heavy chains and light chains are close in expression, as expected since these two have a strong binding affinity (~ 1nM)(61). But clathrin was overexpressed compared to its adaptor proteins by over 3-fold. Functionally, a full triskelia has up to six binding sites for adaptor proteins, but only one needs to be bound to localize it to the membrane. Hence, it is not strictly necessary for the adaptor proteins to be balanced. However, we found that when balanced copy numbers were used instead of observed copy numbers, vesicles formed faster and with fewer components (Fig 7a) Thus the biological copy numbers do not appear optimized for maximum vesicle formation, though they are sufficient to drive vesicle formation.

**Fig 7.**
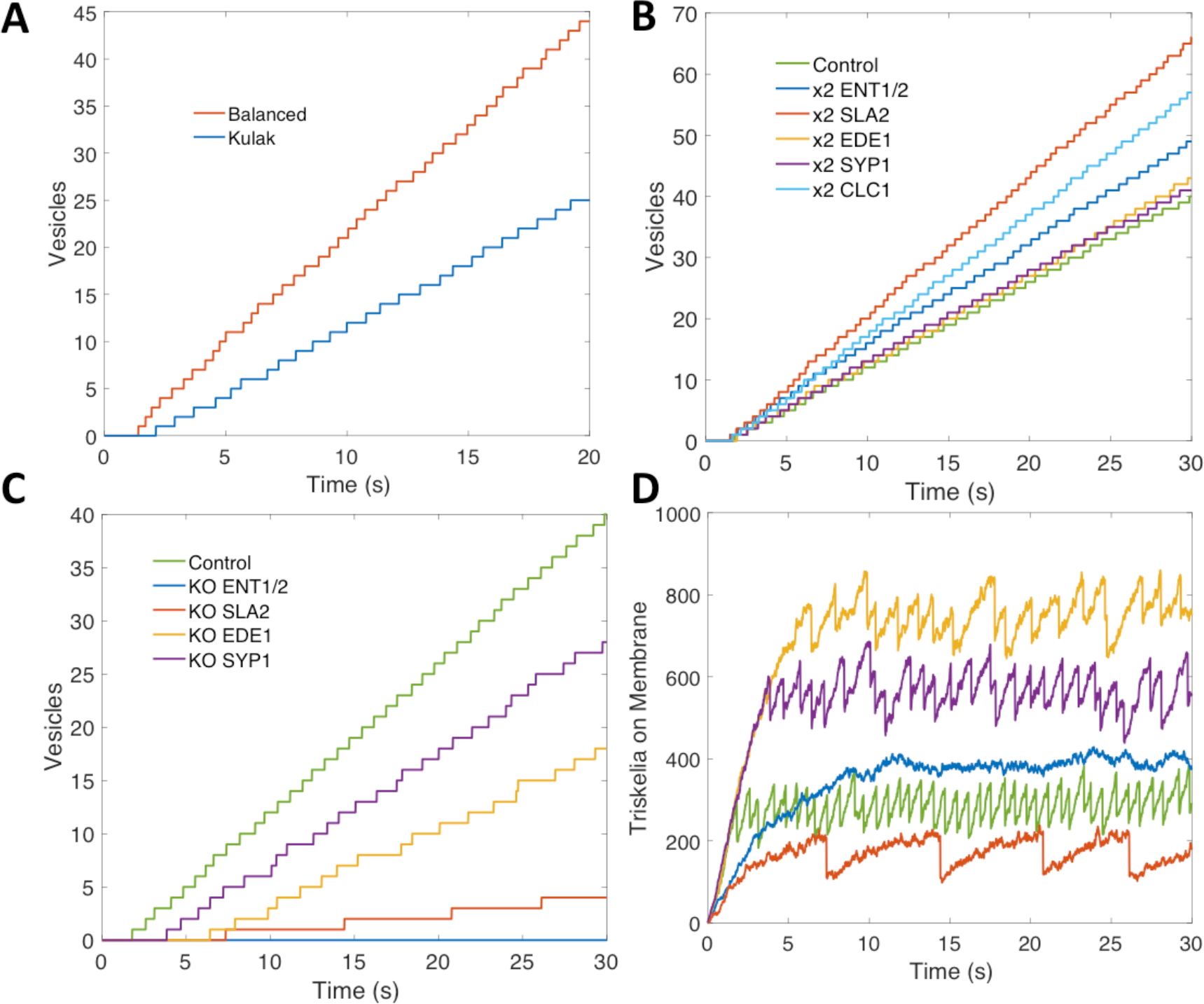
Endocytosis tunable with adaptor proteins. **(A)** Vesicles were formed faster with balanced copy numbers, indicating that the biological copy numbers are not optimized for maximum vesicle formation. **(B)** Adaptor proteins in the network were underexpressed. Vesicle frequency could be increased by doubling their concentrations. **(C,D)** The system is sensitive to adaptor protein knockouts. Knocking out either SYP1 or ENT1/2 nearly halts vesicle formation. SYP1 and EDE1 appear to have an aggregating effect, allowing vesicles to form with less triskelia on the membrane.

Our model assumes these proteins are well-mixed throughout the cytosol, but cells can spatially regulate proteins, altering the local concentration. We simulate this by altering the expression of the adaptor proteins in our model. Knocking out either SLA2 or ENT1/2 pushes the copy numbers even further out-of balance, and nearly halts vesicle formation (Fig 7c,d). Increasing their expression increases vesicle formation because they are below saturation. Decreasing the other adaptor or scaffold proteins also increases imbalance and has a negative effect on the speed of vesicles, although it is less severe. Clathrin-coat assembly is quite sensitive to these membrane-binding protein concentrations because they not only recruit clathrin to the membrane, but they stabilize the triskelion in 2D, where they can then exploit reduced dimensionality to drive binding(69). If clathrin polymerized effectively in solution, far fewer adaptor proteins would be needed to link large clathrin-cages to the membrane surface. We speculate that this sensitivity to the membrane-binding adaptor proteins and their observed underexpression could allow the cell to better tune productive vesicle formation to occur only when enough cargo is localized (70). The adaptor proteins ultimately localize the cargo bound membrane receptors to clathrin-coated sites, a process called cargo loading(71, 72). By increasing or decreasing the local concentration of adaptors, clathrin recruitment can be halted or sped up. With balanced copy numbers, the process is more stable to perturbations in copy numbers, and therefore less efficiently tuned.

Despite the underexpression of adaptor proteins, we observed a very high adaptor to triskelia ratio in completed vesicles (~19). A single triskelion can bind three SLA2 and three ENT1/2 proteins, which can bind three EDE1 and SYP1 proteins, leading to a seeming saturation of 12 adaptors per triskelion. However, most of these proteins can also dimerize with a strong affinity, allowing them to bind to other complexes of adaptor proteins. Our model lacks steric hindrance that would otherwise prevent this high level of aggregation, but nonetheless there is a clear gap in strength between adaptor protein interactions and clathrin interactions (Table 1). These weak clathrin interactions, particularly polymerization (~100 μM)(55), prevent spontaneous cage formation in the cytosol. It is the aggregation of adaptor proteins and localization to the 2D cell membrane that allows cage formation to occur; at least 81% of triskelia were brought to the membrane by adaptor proteins. This suggests another possible reason for overexpression of clathrin: to compensate for lower binding affinity by saturating adaptor proteins.

#### 3.4 Misinteractions have a significant impact for the strong-binding interactions

To determine the overall influence of misinteractions on vesicle formation, and its dependence on protein binding affinity, we added misinteractions at two different strengths (Methods), with an average ratio of K_D,nonspecific_ to K_D,specific_ of 10,000 and 1,000. Despite the weakness of the misinteractions, they decreased the frequency of vesicle formation (Fig 8a,b), though this effect was overall less significant than that of copy number alteration (Fig 7).

**Fig 8.**
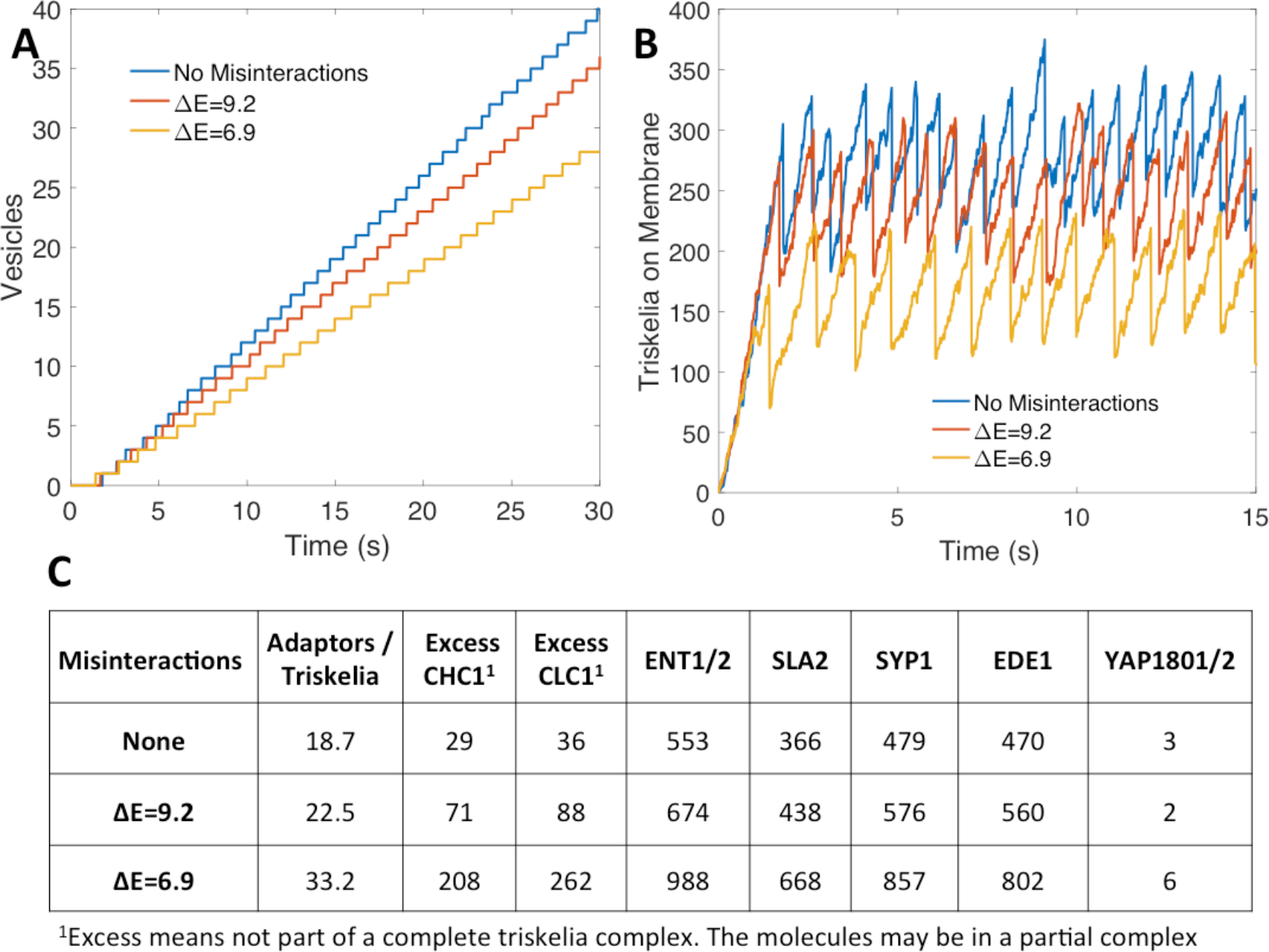
Misinteractions interfere with clathrin recruitment. **(A)** Adding misinteractions to the network decreased vesicle formation and **(B)** interferes with recruitment of triskelia to the membrane. This was caused by aggregates containing too many adaptor proteins, draining them from the cytoplasmic pool. **(C)** Average adaptor proteins in each vesicle. With strong misinteractions, vesicle aggregates contained many adaptors and incomplete triskelia.

In section 2, we found that strong-binding proteins are more sensitive to stoichiometric balance because they are prone to misinteractions. The strongest binders in the network are the Clathrin heavy-chain to light chain interaction (Table 1), and they are both more highly expressed relative to the adaptor partners. Misinteractions dramatically increased the number of both heavy and light chains that were not properly assembled into triskelion (~10 fold), because they became trapped in misinteractions (Fig 8c). For the weaker binding adaptor proteins, the misinteractions increased non-functional aggregation but to a much lower extent, resulting in about 2-fold increase of adaptor proteins in vesicle complexes. Although this 2-fold increase may seem high given the weakness of the misinteractions, it is driven by the localization of these adaptor proteins on the membrane, which concentrates the proteins and promotes binding between any pair of available binding interfaces (69).

Ultimately, misinteractions reduced the frequency of vesicle formation because each vesicle contained a very large aggregate of proteins that drained the cytoplasmic pool of adaptors needed to form new vesicles. The adaptor protein composition is shown in Fig 8c. Without misinteractions, vesicles had an average of 18.7 adaptor proteins per full triskelia, whereas strong misintearctions increased the ratio to 33.2. An interesting consequence of misinteractions is that it initially sped up the formation of the first vesicle, due to the large aggregates assembling on the membrane. However, subsequent vesicles were slower to accumulate than without misinteractions. In contrast, without misinteractions, the speed of initial vesicle formation always correlates with the speed of subsequent vesicles formed.

## Discussion

### Measuring stoichiometric balance in protein-protein networks determines unexpected correlations in protein expression levels

The metric we have developed objectively determines whether a protein is under or overexpressed relative to not only its direct binding partners, but to a larger network including partners of partners. This global evaluation is thus sensitive to the size of the network, but directly captures how the multiple binding interfaces of a protein can control its competition for binding partners. In the interface-resolved CME network, we have shown evidence of imperfect, but statistically significant stoichiometric balance. However, the original 56-protein network was overall unbalanced due to the high overexpression of the actin binding protein cofilin. The size of the network clearly matters, in the small modules, we are statistically out-of-balance, but on a larger scale, still in balance. Outliers are emphasized in smaller networks. At the same time, leaving out additional partners can provide some explanation for the observed imbalance. Imbalance may also indicate possible missing interactions in the network. Despite the simplicity of our metric, our method was still able to highlight both correlated concentrations and proteins that violate balance for functional reasons, such as the kinase PRK1. Furthermore, the observed balances can suggest possible mechanisms of assembly, for example, that can then be studied using kinetic modeling, as we did here. What our results emphasize is that correlations are highly important: functionality can be obliterated with significant imbalance, and misinteractions can also be overwhelming with significant imbalance.

Although we only applied our stoichiometric balance analysis to the 56 protein CME network, two smaller modules of this network, and the 127-protein ErbB network, these networks are significantly larger than the obligate complexes previous studied for copy number balance(5, 6). Our networks also contain a much larger variety of binding interaction strengths and competitive and non-competitive interactions. As we showed above, balance depended on the protein network’s underlying IIN. While it would be beneficial to repeat this analysis on a larger network, there is a paucity of manually curated IINs in the literature. There are various larger automatically constructed IINs, constructed with homology modeling(73, 74), but our previous work found these automatic IINs suffer from various inaccuracies and differ significantly from manually curated IINs in topology(41).

### Limitations of measuring stoichiometric balance for larger PPINs

Our metric for evaluating stoichiometric balance only accounts for the binding interface network structure and observed copy numbers. A missing feature of our stoichiometric balance metric is that proteins within a network can be expressed with both spatial and temporal variation. For a small binding network this is not a major concern, since proteins in the same complex tend to be co-expressed(75) and co-localized so they may bind. But as network size is scaled up, the probability of all proteins being equally present reduces. Such temporal and spatial variations could be taken into account in the construction of the network, leaving out proteins that are not functional at the same time.

A natural extension to our measure of stoichiometric balance would be to also account for binding affinities of interactions in addition to the binding interface network structure and observed copy numbers. Our results here and previous studies(19) indicate that balance should be more tightly constrained for strong binding proteins. However, one benefit to leaving affinities out of the measurement is that biochemical data is in even more limited availability than binding interface data. Our existing metric can thus be much more easily applied to a variety of networks. Furthermore, by picking out highly correlated expression levels, our method can then indicate which interactions might be quite strong, or vice-versa, which may be transient or weak.

### Noise and variability in experimental copy number measurements can limit observed balance

In this study we used yeast copy numbers from Kulak et al. because it was the most comprehensive. The other three studies we used for comparison did not cover all 56 proteins in our network. However, for the proteins we could compare, we found significant discrepancies between relative abundances. Light chains are weakly expressed in other studies, for example(1, 47, 48) A few possible reasons for this exist. The first is that fluorescence data is inherently noisy. Experimentalists must deal with background noise, interference with protein localization due to the large fluorescent tags, and cross interactions with other proteins(76). The second is that cell lines can accrue mutations over time that decrease or increase gene expression, a phenomenon observed with HeLa cells(77). Finally, cells may alter gene expression for regulatory reasons, so the environment in which cells are grown may alter gene expression.

### Perfect balance is not observed, even if it would improve both misinteractions or the functional outcome of the protein network

We do not expect the cell to perfectly optimize the yield of all of its many assemblies. Each network we have evaluated here is ultimately part of a larger, global cellular network. Perfectly optimizing isolated, local modules does not appear to be a significant pressure for the cell, particularly when a sufficient balance, such as we observe for the vesicle-forming module, maintains functionality. Correlations in copy numbers are nonetheless often significant relative to randomly assigned copy numbers.

We found that copy number imbalance can lead to misinteractions and the features of biological IINs (power-law-like degree distribution, square and hub motifs, sparseness) typically have less misinteractions under balance copy numbers but more misinteractions under imbalance. These networks thus should require more tightly controlled balance to avoid misinteractions. But misinteractions are of course not the only pressure on copy numbers. For multi-protein assembly in an obligate complex (ARP2/3) and in a minimal model of vesicle formation for CME, we found that the functional cost of imbalance was dominated more by its impact on determining specific functional complexes than avoiding misinteractions. Nonetheless, the fact that misinteractions can decrease vesicle formation, by sequestering away adaptor proteins into large aggregates, shows that misinteractions are worse than simply having an excess of free proteins. If this result can be generalized, it may have important implications for mechanistic modeling of biological systems, as misinteractions or system error is rarely taken into account.

### Observed imbalances in the non-equilibrium vesicle forming module could provide benefits to assembling cargo-selective vesicles

Although the functional effects of copy number balance are usually discussed in the context of number of complete complexes at equilibrium, we have shown that nonequilibrium dynamics can be affected as well. While the clathrin heavy chains and light chains were balanced with each other, they were overexpressed compared to their adaptor proteins, and this limited the frequency of vesicle formation. Although we found that perfectly balanced copy numbers therefore improved vesicle formation frequency compared to observed copy numbers, we speculate that specific imbalances could still be selected for evolutionarily. There are various possible reasons for this imbalance: the function of endocytosis is cargo uptake, and there is a cargo loading process before endocytosis occurs.(71, 72) Hence to maximize function, controlled endocytosis around high-cargo areas of the membrane may be preferably to frequent, spontaneous endocytosis, and the adaptor proteins can serve as an intentional bottleneck in the process. However, the observed underexpression could also be because there are other adaptor proteins not included in our model, or because clathrin interactions have weaker affinities than interactions between adaptor proteins and must saturate them.

Finally, the predictions of our minimal vesicle-forming model are ultimately limited by the approximations we made to simulate the clathrin coat assembly and vesicle formation. Our model vesicles formed about 10 times faster than is observed *in vivo*. To fully capture the dynamics of this complex process, an ideal model would include all the proteins in our CME network (Fig 1), and include both the known biochemistry of binding interactions and the physics and biomechanics of membrane bending and scission. In yeast, the cytoskeleton is needed to help induce membrane budding, after which energy-consuming proteins such as dynamin scission off the vesicle from the plasma membrane for transport into the cell (72, 78). However, such a modeling approach does not exist, due to the computational limitations of simulating such large complexes and membrane remodeling, and the lack of biochemical data.

Based on the model we did construct, however, there are some more specific limitations. The first is that while rule-based modeling is a convenient way to model complex formation, some theoretical aggregates may be impossible due to steric hindrance. Our model predicted that a vesicle of 100 triskelia could contain ~1900 additional proteins. Assuming each vesicle is a sphere with 100nm diameter, the allowable surface area per adaptor/scaffold protein would only be ~17nm^2^, which is too small to accommodate the excluded volume of the large, disordered regions of proteins such as ENT1 and 2{Busch, 2015 #304}. Second, we did not include cooperatively in our model. Molecules localized in the same aggregate do not interact at a faster rate in conventional rule-based modeling. Clathrin triskelia weakly polymerize, as noted above, but the aggregation effect of the adaptor proteins - especially the SYP1/EDE1 complex - localizes triskelia close together, allowing them to bind strongly. In future work we will consider effects of cooperativity on assembly, as well as construct more detailed spatial and structural models of the vesicle forming process.

## Methods

### Defining stoichiometric balance in a PPIN with interfaces resolved

A stoichiometrically balanced network has the copy numbers of each interface matched to the copy numbers of all pairwise complexes it participates in. Balanced copy numbers are obtained by assigning a number of desired complexes to each edge in the interface binding network. The balanced copy numbers of each interface can then be calculated from the equation:

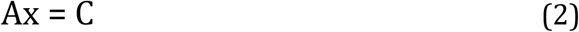

Where “A” is a binary matrix with N_int_ rows (one for each interface) and M_edge_ columns (one for each pairwise interaction). A_i,j_ =1 if the interface *i* is used in the interaction *j*, or 2 if a self-interaction, and 0 otherwise. “x” is the vector of desired pairwise complexes (M_edge_ × 1), and “C” is the number of interface copy numbers (N_int_ × 1). In Fig 9 we illustrate this procedure for a small toy network.

**Fig 9.**
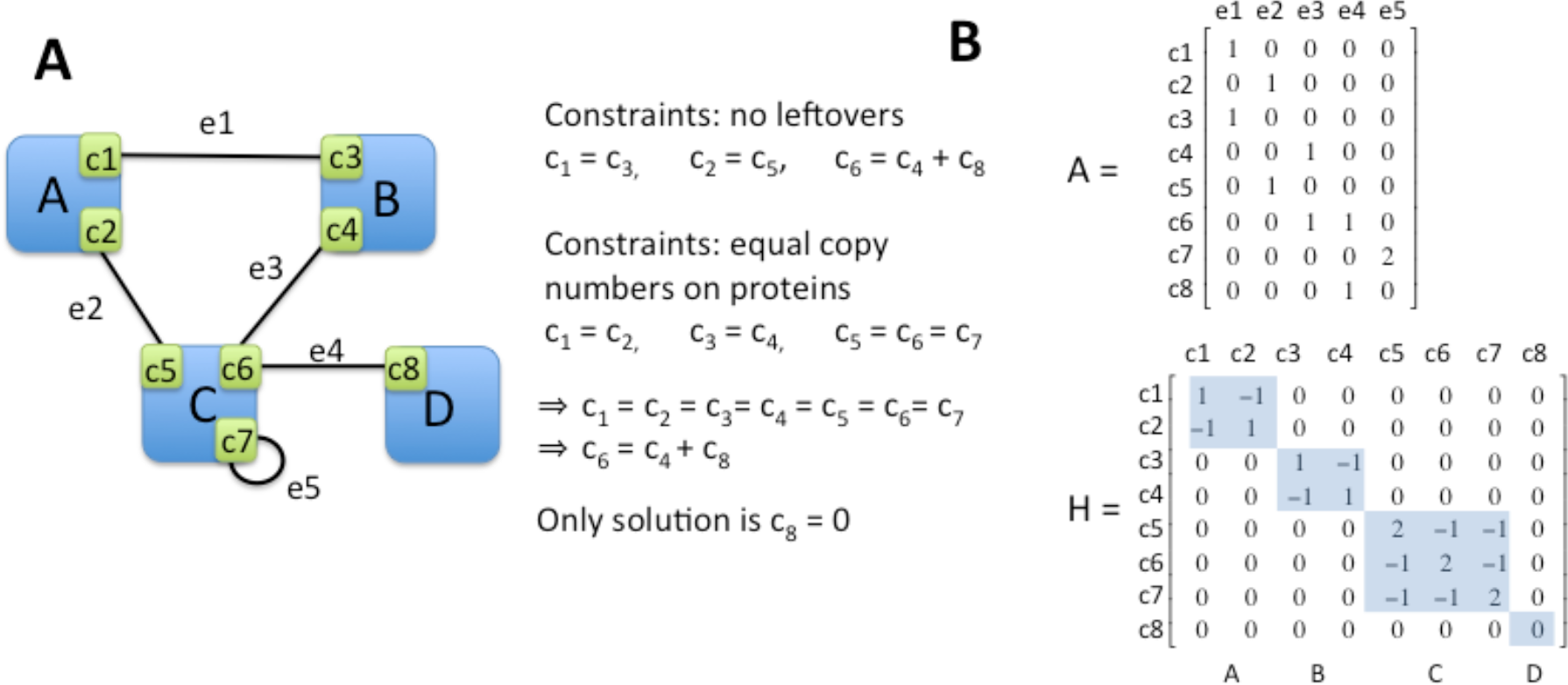
Example network for calculation of stoichiometric balance. **(A)** An example of an interface-resolved protein network that has no nontrivial balanced solution when all constraints are applied. **(B)** To solve for balanced copy numbers, the network topology is encoded in the “A” and “H” matrices illustrated here for the network in (a).

If desired pairwise complexes, x, is specified, interface copy numbers, C, can directly be solved for using Eq. 1, but if interface copy numbers, C, are specified, x will not, in general, have an exact or nontrivial solution unless C is balanced. This is because all entries of x must be >0 or some other minimum value, as negative copies cannot exist. This produces a hard constraint on x. Given a vector C, an optimal solution to x must be solved for using quadratic programming rather than linear least-squares.

Our goal is to select for an optimal x given an input set of copy numbers “C_0_”. This is a soft constraint on the optimal x, because the input C_0_ may not be balanced. Once an optimal x is found, forward solving Eq. 1 will in general not perfectly recover C_0_. C_0_ can constrain all interfaces or a subset of them. To constrain a protein is to constrain all interfaces on it. We introduce a third constraint on the optimal x: the copy numbers of interfaces on the same protein should be equal. This often makes nontrivial solutions impossible (Fig 9), so it is also a soft constraint. Combining all of these constraints, the optimal desired number of complexes “x” can be found by minimizing the equation:

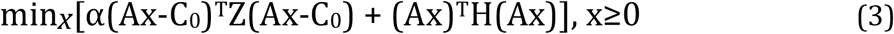

Where each variable is defined as follows:

A: N_int_ x M_edge_ matrix defining which interfaces are used in which interaction, i.e. pairwise complex.
x: M_edge_ x 1 vector of desired pairwise complex copy numbers
C_0_: N_int_ x 1 vector of constrained copy numbers.
Z: N_int_ x N_int_ diagonal matrix that selects which interfaces are constrained. Entries = 1 if the interface is constrained and =0 otherwise. If all interfaces are constrained, Z equals the identity matrix.
H: N_int_ x N_int_ permutated block diagonal matrix with positive and negative entries such that H*C=0 if interfaces on the same protein have equal copy numbers. Each block corresponds to a protein (Fig 9).
α: 1x1 scaling parameter which determines the relative weight of the C_0_ soft constraint vs the equal interfaces soft constraint.

For any vector x, Eq. 2 produces a positive scalar value. The equation was minimized using the OOQP (object-oriented quadratic programming) 0.99.26 package for C++(79). Quadratic programming is necessary due to the constraint of x≥0. Eq. 2 can be converted into a quadratic equation of the form

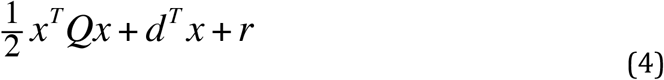

Using

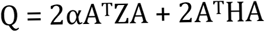

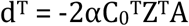

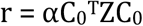

“r” can be ignored by the solver when minimizing the equation since it is a constant term.

Once x_min_ is found via Eq. 3, the optimized interface copy numbers can obtained by forward solving A*x_min_ = C_balanced_. Interfaces on the same protein will not necessarily have equal copy numbers due to the competing constraints of Eq. 2. Distance from C_0_ to C_balanced_ was used as a metric to determine relative balance (see below). We can assign a single copy number to each protein by averaging over all interface copy numbers on that protein to give P_balanced_, a vector of protein copy numbers. These values were only used when visualizing which proteins were over or underexpressed in the networks and highlighting these protein copy numbers (Fig 2e).

### Biological protein copy numbers

All copy numbers are collected in S2 Table. For the yeast CME network, C_0_ was used to constrain all 56 proteins (Z=Identity matrix) because copy numbers from Kulak et al. were available(2). For the ErbB signaling network, only 115 out of 127 proteins with available expression level data were constrained. 1oo of these proteins were constrained with HeLa copy number estimations from Kulak et al. (2), while estimated copy numbers for 15 additional proteins were added from four additional studies(19, 49–51), leaving 12 proteins with unknown expression data.

### Measuring the degree of stoichiometric balance in observed concentrations

Using the optimized copy numbers, C_balanced_, we can then ask, how close are the original, biologically observed copy numbers to these optimally balanced values? If the original copy numbers are already perfectly balanced, then they will match the optimal copy numbers. If they are imperfect, than the two distributions will differ. We use two metrics to quantify the distance between the observed and optimized concentrations: chi-square distance (CSD)

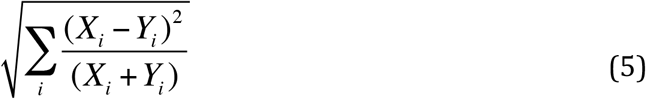

and Jensen-Shannon Distance (JSD) after converting both vectors (X and Y) to distributions (x and y)

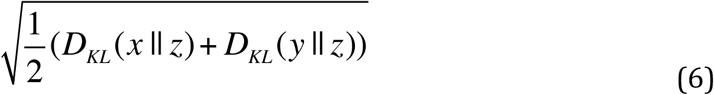

Where z=(x+y)/2 and D_KL_ is the Kullback-Leibler divergence

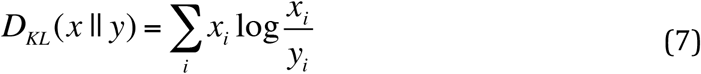

For cases where Z≠I (i.e. not all interfaces were constrained) only distance between constrained interfaces was measured.

### Small network motifs

Binding for the five 3- or 4-node network motifs; triangle, chain, square, 4-node hub, and flag; was simulated using the Gillespie algorithm(45). Besides the specific binary interactions, nonspecific interactions were allowed at a strength determined by an “energy gap” between binding energies, though in practice we defined the ratio nonspecific K_D_ to specific K_D_ by factors of 10. This corresponded to a linear difference in free energies via the equations:

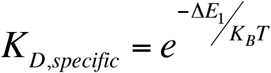

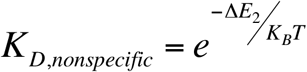

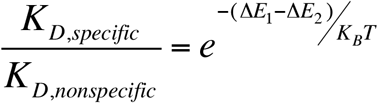

The networks were simulated under various initial concentrations. The steady-state ratio of Eq. 1 was recorded, where N_nonspecific_ is the number of nonspecific binary complexes, N_specific_ is the number of specific binary complexes, and N_free_ is the number of free proteins. Ratios were averaged across 5,000 runs.

To generate surface plots, two proteins were chosen to be variable while the remaining proteins were given fixed copy numbers. Because the flag motif produced asymmetric plots, two different choices of variable proteins were used. (S3 Fig) Surface plots were generated using Matlab.

We calculated sensitivity by determining the principal component of the surface plot data (i.e. the vector of greatest variance) and measuring the percent change in ratio from the optimum along this vector. For better comparison, we normalized distance along the surface plots via dividing the abundance of the variable proteins by the abundance of the fixed proteins.

### Analysis of complex IIN topologies

For the large network analysis we used the 500 networks from Johnson, *J Phys Chem B* 2013(27). 25 sets of 10 networks each were randomly generated using two parameters: number of nodes (90, 110, 125, 150, 200), keeping the number of edges fixed at 150; and the preferential attachment exponent “γ” from Goh, 2001(80). γ=0 corresponds to a binomial, Erdos-Renyi network, whereas γ=1 corresponds to a power-law or “scale-free” network. Values of 0, 0.2, 0.4, 0.6, and 0.8 were used. Finally, a local topology optimization algorithm that decreased the frequency of chain and triangle motifs and increased hub motifs was applied to each network, for 500 networks in total. All networks assume competitive (binary) binding.

Rather than assign an arbitrary specific and nonspecific K_D_ for the networks, we used the relative binding energies determined for each network in the source paper. This was determined by a physics-based Monte Carlo optimization scheme of amino acid residues, as described in Johnson, 2011(23). The minimum energy gap between specific and nonspecific interactions could be measured as a relative metric of the network’s propensity for misinteractions. Because the binding strengths were relative, we could alter the average binding strength to determine the effects on misinteractions. This was varied between 7 values of 1 nM to 1 mM, using factors of 10. Finally, to obtain results more comparable to the simple networks, we also ran simulations where each specific interaction had K_D_=100 nM and each nonspecific interaction had K_D_=100 μM.

Networks were simulated to steady state using the Gillespie algorithm(45) under five differing sets of copy numbers (CNs) for free proteins: equal CNs for each protein, random CNs sampled from a yeast protein concentration distribution (performed 20 times) and three forms of balanced CNs using the network architecture. Any set of CNs without leftovers - i.e. having exactly enough proteins to create a certain number of specific complexes - is considered “balanced”, and thus there are infinite solutions. The first balanced set assumed an equal number of each type of specific complex, which results in protein CNs proportional to the protein’s number of partners. The remaining balanced CNs were determined by finding “x” to minimize a simplified form of Eq. 2:

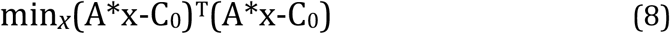

Here there is only one interface on each protein, and all the proteins are constrained, so there is no need for a Z matrix, the α scaling parameter, or the second term. C_0_ is either equal copy numbers or randomly sampled copy numbers.

After x_min_ is found via quadratic programming (see above), the balanced CNs are obtained by forward solving C_balanced_ = A*x_min_.

To measure nonspecific complex formation, a modified ratio was used:

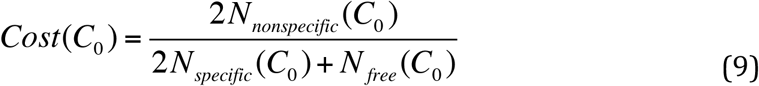

to compare total individual proteins in each bound or unbound state, rather than number of unbound or bound states. To measure sensitivity, the ratio under unbalanced CNs (C_0_) divided by the ratio under balanced CNs (C_balanced_) was calculated. A higher ratio indicates higher sensitivity to CN balancing.

### ARP2/3 Complex

The kinetic model was simulated using the stochastic simulation method (the Gillespie algorithm). Binding interactions were encoded via the rule-based language BioNetGen and simulated via the Network Free Simulation (NFSim) software (46). Trimer cooperativity was modeled by increasing the rate of the third reaction if three members of a correct trimer were held together by two reactions. For example, if A is bound to B is bound to C, and a binding between A and C is possible, that reaction rate was set to be arbitrarily high. Reaction rates were arbitrary, but interactions with the core subunit ARC19 were set to be ~10 fold stronger than interactions between periphery subunits, as this increased yield. Yield was measured via the equation

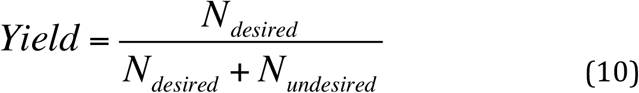

Where N_desired_ is the number of *proteins* in complete complexes (equal to seven times the number of complex complexes) and N_undesired_ is the number of proteins in incomplete or misbound complexes. Completely free proteins were ignored.

### Simulating clathrin recruitment to the membrane

A subnetwork of nine proteins - clathrin heavy chain (CHC1), clathrin light chain (CLC1), SLA2, ENT1/2, EDE1, SYP1, and YAP1801/2 - was defined based on known binding interactions (Table 1). Because the existence of multiple interfaces, allowing noncompetitive binding, results in a large number of possible species we simulated our model using the Network Free Simulator (NFSim)(46). Binding dissociation constants were obtained from the literature, including for protein-lipid binding. For simplicity, the heavy chains were already assumed to be in trimer form, and ENT1/2 was combined into a single protein as the binding partners were the same. Binding constants were pulled from the literature. (Table 1)

The cell membrane and the cell cytoplasm function as different compartments with different volumes, but NFSim is not integrated with BioNetGen’s compartment language. We bypassed this problem by doubling the number of rules: besides the main rule for each reaction, an additional rule stated that if both proteins are on the cell membrane then the k_on_ rate should be increased according to the membrane volume. Cell membrane ‘volume’ was determined by multiplying the membrane surface area by a factor 2σ=2 nm to capture the change in binding affinities between 3D and 2D (see SI Text).

Since our primary goal was to measure clathrin recruitment to the membrane, any complex on the membrane with at least 100 triskelia (a complex of three CHC1 and three CLC1) was considered a “vesicle” and deleted at a high rate k_dump_. Proteins in the vesicle were then added back to the cytoplasmic pool at a rate k_recyc_, which was set to be equal to k_dump_ to indicate fast recycling. However, we clarify that even fast recycling is not instantaneous, and that proteins are added back one at a time rather than all at once. Fast vesicle formation thus could still drain the pool of adaptor proteins.

Misinteraction strengths were determined by calculating the geometric mean of the dissociation constants of each interface, as this provided a K_D_ based on the arithmetic mean of the binding energies.

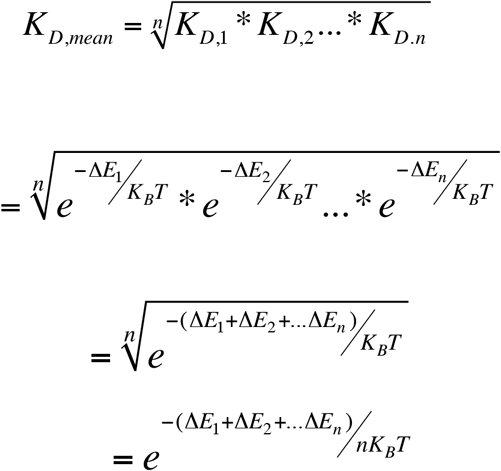

The K_D_ of a misinteraction between two interfaces was set to be:

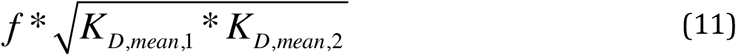

where f=10,000 (weak misinteractions, corresponding to an energy gap of ~9.21) or 1,000 (stronger misinteractions, energy gap of ~6.91)

Network maps were generated using Cytoscape(81) and RuleBender(82). Plots were generated in MATLAB.

## Acknowledgements

Research reported in this publication was supported by the National Institute of General Medical Sciences of the NIH under Award No. R00GM098371 to M.E.J. We gratefully acknowledge computational resources from Maryland Advanced Computing Cluster (MARCC) and the Hopkins High Performance Cluster (HHPC). We thank Daisy Duan for helping to collect rate parameters and the Johnson lab members for helpful feedback on the manuscript.

## Supporting Information

### SI.pdf

**S1 Text. Notes on the vesicle forming module**

**S1 Table. Model parameters with notes**

**S1 Fig. Effects of the α parameter on interface copy number noise**

**S2 Fig. Upstream proteins in the ErbB network are underexpressed**

**S3 Fig. Misinteraction frequency in the small networks**

**S4 Fig. Effects of optimized local topology on misinteractions**

**Fig. ARP2/3 complex has higher yield under balanced copy numbers.**

**S2 Table (Excel) S2Table_ProteinCopyNumbers.xlsx** Copy numbers used for the CME and ErbB Protein networks

